# Partial loss of succinate dehydrogenase reduces high red cell distribution width and promotes healthy survival in chronically hypoxic mice

**DOI:** 10.1101/2021.05.18.444547

**Authors:** Bora E. Baysal, Debra Tabaczynski, Leslie Curtin, Mukund Seshadri, Sandra Sexton

**Author notes:** To whom correspondence should be addressed: Bora E. Baysal MD, PhD, Department of Pathology and Laboratory Medicine, Roswell Park Comprehensive Cancer Center, Elm and Carlton Streets, Buffalo, NY 14263, Lab.

## Abstract

Increased red cell distribution width (RDW), which measures erythrocyte size variability (anisocytosis), has been linked to early mortality in many diseases and normal aged population through unknown mechanisms. Hypoxia has been proposed to increase both RDW and mortality. However, experimental evidence, especially in animal models, is lacking. Here, we show that chronic hypobaric hypoxia (~10% O_2_) increases erythrocyte numbers, hemoglobin and RDW, while reducing longevity in male mice. Compound heterozygous knockout (chKO) mutations in succinate dehydrogenase (Sdh; mitochondrial complex II) genes *Sdhb, Sdhc* and *Sdhd* reduce high RDW and immature reticulocyte fraction, and increase healthy lifespan in chronic hypoxia. Hemoglobin and erythrocyte numbers in hypoxia do not show statistically significant differences between Sdh chKO and WT mice. These results identify a mitochondrial mechanism regulating both RDW and organismal adaptation to chronic hypoxia, and suggest SDH as a potential therapeutic target to reduce high RDW-associated clinical mortality.

## Introduction

Erythrocyte anisocytosis refers to increased variation in red blood cell (RBC) size, and is measured by red cell distribution width (RDW) in routine complete blood count (CBC) analysis. RDW is often reported as a coefficient of variation (RDW-CV) of erythrocyte mean corpuscular volume (MCV) in RBC volume distribution curve. RDW-CV is calculated by dividing the standard deviation (SD) by MCV, multiplied by 100. RDW-SD is a direct measure of anisocytosis that reports the MCV variation at 20% frequency level [1]. RDW increases in healthy aging population [2].

RDW along with MCV was traditionally used in the differential diagnosis of anemia. In recent years, however, high RDW has been associated with increased mortality in acute and chronic diseases as well as in middle-aged and older individuals without disease [3, 4]. The association has been observed in a growing list of clinical conditions including heart failure [5, 6], myocardial infarction [7], peripheral artery disease [8], cancer [9], pulmonary hypertension [10], acute pulmonary embolism [11], community-acquired pneumonia [12, 13], SARS-CoV-2 infection [14], chronic obstructive pulmonary disease [15, 16], acute respiratory distress syndrome [17], acute cerebral infarction and stroke [18, 19], intensive care unit (ICU) and trauma patients [20–22], hip fracture [23], sepsis and septic shock [24], gram-negative bacteremia [25], acute pancreatitis [26], hemodialysis [27], and kidney transplant receivers [28]. The underlying mechanism(s) for this association is unknown.

Although anisocytosis is a physiologic response to anemia, the association with mortality in non-anemic individuals remains significant [29–34], and becomes even stronger than seen in anemic individuals in meta-analysis [35]. A correlation between inflammatory markers and anisocytosis has been documented, which raises the hypothesis that RDW effect on mortality may be mediated by systemic inflammation [36]. However, the association of anisocytosis with mortality and disease remains statistically significant even in subjects with low inflammatory marker CRP levels [4, 37, 38]. It has been hypothesized that RBCs with increased size variation may have reduced deformability that impairs micro-circulatory blood flow, though contrasting results were reported on the impact of increased RDW on RBC deformability [39, 40]. Furthermore, certain anemias cause anisocytosis without significantly increasing the mortality risk. For example, dietary iron deficiency anemia is attributed to ~0.08 deaths per 100,000 [41]. These considerations collectively suggest that the mortality risk associated with anisocytosis cannot be readily explained by anemia, inflammation or RBC physicochemical characteristics.

Increased anisocytosis may reflect a fundamental cellular pathology that predisposes to mortality regardless of the specific clinical condition. Yčas et al. analyzed over 2 million medical claims and concluded that RDW indicates systemic hypoxic load, especially in pulmonary and cardiac conditions [42]. High RDW correlates with severity and poor survival in COPD [15, 16, 43, 44] as well as with lung function in normal subjects [45]. A recent analysis of RDW in 121,530 non-anemic individuals with a medical condition revealed the strongest associations with pulmonary hypertension, chronic pulmonary heart disease and congestive heart failure, which all have a pathophysiologic link to hypoxia [46]. Similarly both hypoxemia and high RDW have been linked to mortality risk in COVID-19 patients [14, 47]. Hypoxia triggers the production of RBC precursor reticulocytes from bone marrow through the operation of PhD-HIF pathway that regulates erythropoietin production[48]. Since reticulocytes are larger than mature RBCs, anisocytosis ensues. Thus, systemic hypoxia appears to be a biologically plausible factor that might explain the association between anisocytosis and mortality. However, experimental evidence for this hypothesis is lacking.

In this study, we report on the impact of chronic hypobaric hypoxia on RBC parameters and survival in Sdh heterozygous and wild-type control male mice. In humans, heterozygous germline SDH subunit mutations predispose to paraganglioma (PGL) and pheochromocytoma tumors [49]. Hereditary PGL tumors caused by *SDHD* mutations often develop in the carotid body (CB) in neck [50], and mimic the sporadic CB paragangliomas caused by chronic hypoxic stimulation of high altitudes [51]. Higher altitude increases the severity of hereditary PGL tumors [52, 53]. Gene expression profiling studies in SDH PGLs show persistent activation of hypoxia-induced genes in normoxic conditions (pseudo-hypoxia)[54]. These results collectively suggest that SDH mutations predispose to PGL tumors by constitutively activating the hypoxia-sensing/signaling pathways in paraganglionic tissues.

Sdh mouse models show that while homozygous deficiency of a subunit is incompatible with normal life and development, heterozygous mutations do not cause paraganglioma tumors [55]. Here, we present evidence that chronic hypoxic stimulation also fails to develop paraganglioma tumors in Sdh mice. We find that mice with partial Sdh deficiency show reduced RDW and increased survival relative to control mice in chronic hypoxia, revealing an unexpected mechanism contributing to the association between high RDW and mortality.

## Methods

### Sdh knockout mice

The experimental mice were derived by crossing three previously described original strains each containing a heterozygous knockout mutation in a distinct Sdh subunit (*Sdhb, Sdhc* or *Sdhd*). Sdhb and Sdhc heterozygous KO mice were created in The Jackson Laboratory (Bar Harbor, Maine) in B6/129P2 background [56]. The original strains are described as: B6.129P2-Sdhb<Gt(AP0532)Wtsi>/Cx and B6.129P2-Sdhc<Gt(BA0521)Wtsi>/Cx. *Sdhd* knockout mouse [57] was re-derived into C57BL/6J background at RPCCC transgenic facilities using frozen sperm (mfd Diagnostics, Germany). As previously reported, homozygous mutations in any subunit are non-viable, but compound *Sdhb/Sdhc* double heterozygous and *Sdhb/Shdc/Sdhd* triple heterozygous KO mice are viable [56]. Since each gene is located on a different mouse chromosome, the KO alleles segregate independently and give the expected numbers of each viable genotype upon crossing Sdh hKO mice. Control WT mice were also derived from crosses of Sdh hKO mice. WT controls were either littermates or from closely related litters. If WTs Genotyping was performed in tail tips at RPCCC transgenic facility as described [56]. Genotypes of the mice in hypoxia chamber are confirmed by repeat testing.

### Hypoxia exposure

Mice were exposed to chronic hypobaric hypoxia in a custom-made hypoxia chamber (Case Western Reserve University Design Fabrication Center, Cleveland, OH) that operates via house vacuum and accommodates 2 standard mice cages, as previously described [56]. Mice were initially subjected mild hypoxia (~15%) for ~1 week for acclimatization. For chronic exposure. the oxygen concentration was ~10% with a range of 9–11%, due to house vacuum oscillations. Oxygen percentage is continuously monitored by an O_2_ sensor. Hypoxia exposure experiments involved 5 compound heterozygous and 5 WT control mice, with each genotypic group placed in a different cage after ~ 10 weeks of age. Mice were daily observed, and briefly removed from the chamber twice a week for cage cleaning.

Mice remained in hypoxia chamber until spontaneous death or the development of morbid conditions, as assessed during cage cleaning, that required euthanasia in accordance with Roswell animal care guidelines and the approved IACUC protocol. Examples of morbid conditions included limited or absent movement, hunched posture, labored breathing, sunken eyes, shaking and development of rectal prolapse. All decisions for euthanasia due to morbid status were made in accordance with approved IACUC guidelines. Organs were grossly examined during necropsy. Tissues were collected upon spontaneous death and euthanasia.

### Peripheral blood collection

Periodically, body weights were measured and blood was collected for CBC analysis. Blood (~0.2 mL) is collected into EDTA tubes by retro-orbital bleeding at baseline and subsequent time points. Alternate eyes were used for a maximum of 2 times per eye. Additional bleeding was performed by mandibular venipuncture. Complete blood counts were analyzed via automated cell counters Hemagen HC5 (Group 1) or ProCyte Dx (Groups 2 and 3) hematology analyzers through Roswell Laboratory Animal Shared Resources. Certain RBC parameters such as RDW-SD and IRF were not reported by Hemagen HC5 counter. The RBC parameter values examined in this study are direct outputs of the analyzers except 1SD-RDW, which is derived as (RDW-CVXMCV)/100.

### Statistical analysis

Statistical analysis are performed by GraphPad Prism (Versions 7.03 and 9.1.0). CBC output values are first entered to Microsoft Excel and then imported into GraphPad. A few extreme outlier values were removed using the most stringent criteria of GraphPad’s ROUT method, that removes 0.1% (Q=0.1%) of the data obtained at a time point. Statistical analysis and graphic presentations were performed using GraphPad Prism. The comparisons of CBC values over time between Sdh chKO and WT control were performed by 2-way ANOVA test by using data from all available time points, including the baseline normoxic values, within each group. The independent variables are time and genotype. When the genotype comparison was made in normoxia or hypoxia, data from all time points from all three groups were combined. The survival differences were calculated by Kaplan-Meier method, where the outcome is time until death or the development of morbid conditions that required euthanasia according to institutional guidelines.

## Results

### Sdh heterozygous knockout mice do not develop tumors under chronic hypoxia

The development of PGL or other tumors was prospectively examined in 3 sequentially tested groups of male mice exposed to lifelong hypoxia (~10% O_2_). Group 1 had Sdh double hKO of *Sdhb* and *Sdhc*, whereas Groups 2 and 3 had Sdh triple hKO of *Sdhb, Sdhc* and *Sdhd* (Table 1). The initial goal was to determine whether compound heterozygosity in Sdh predisposes to PGL tumor development under chronic hypoxia. MRI analysis of 1 chKO Sdh bc (#145) and 1 WT control (#197) mouse after ~7 months of chronic hypoxia exposure showed no MRI evidence of tumor development in either genotype (**Supp Fig. 1**). Gross and microscopic examination of the hypoxia-exposed mice showed no evidence of tumor development or vascular pathology including intimal thickening or plexiform lesions in lung or pheochromocytoma development in adrenal gland (**Supp. Fig. 2**). Thus, although hypobaric hypoxia of high altitudes promotes development of sporadic paragangliomas in humans, we find no evidence of Sdh-related PGL tumor development in mice both in normoxia and hypoxia.

**Table 1.** Longevity of mice under chronic hypobaric hypoxia.

| **Group no.**<br>**(start date)** | **Mouse**<br>**ID** | **Genotype** | **Survival**<br>**time**<br>**(days)** | **Manner of**<br>**death** | **Euthanasia**<br>**indication** |
| --- | --- | --- | --- | --- | --- |
| 1 (03/25/13) | M194 | WT | 461 | Spontaneous | - |
|  | M195 | WT | 441 | Spontaneous | - |
|  | M196 | WT | 211 | Spontaneous | - |
|  | M197 | WT | 211 | Euthanasia | Elective for tumor<br>evaluation |
|  | M198 | WT | 420 | Spontaneous | - |
|  | M145 | Sdh bc | 211 | Euthanasia | Elective for tumor<br>evaluation |
|  | M146 | Sdh bc | 494 | Euthanasia | Moribund |
|  | M147 | Sdh bc | 514 | Spontaneous | - |
|  | M148 | Sdh bc | 256 | Spontaneous | - |
|  | M185 | Sdh bc | 503 | Spontaneous | - |
| 2 (06/09/15) | M598 | WT | 476 | Euthanasia | Moribund<sup>#</sup> |
|  | M599 | WT | 490 | Spontaneous | - |
|  | M600 | WT | 486 | Euthanasia | Moribund<sup>#</sup> |
|  | M601 | WT | 454 | Spontaneous | - |
|  | M602 | WT | 456 | Spontaneous | - |
|  | M515 | Sdh bcd | 527 | Euthanasia | Elective<sup>^</sup> |
|  | M518 | Sdh bcd | 495 | Spontaneous | - |
|  | M527 | Sdh bcd | 495 | Spontaneous | - |
|  | M531 | Sdh bcd | 524 | Spontaneous | - |
|  | M536 | Sdh bcd | 527 | Euthanasia | Elective<sup>^</sup> |
| 3 (08/18/17)* | M773 | WT | 371 | Euthanasia | Weight loss, rectal<br>prolapse |
|  | M803 | WT | 222 | Euthanasia | Rectal prolapse |

^M515 and M536 were the last surviving mice in group 2, and euthanized electively. # M598 and M600 showed limited spontaneous mobility (videos available). \*The experiment with Group 3 stopped on day 446, when there were 3 WT and 5 Sdh bcd alive, due to hypoxia chamber failure. These and the four electively euthanized mice are censored in Kaplan Meier survival analysis.

### Sdh heterozygous knockout mice survive longer under chronic hypoxia

We observed mice until spontaneous death or development of morbidities that require euthanasia to increase the possible risk of hypoxia-induced paraganglioma development. Healthy lifespans of mice in chronic hypoxia (median 503 days for Sdh chKO and 456 days for WT control) were substantially lower than that of the parental B6 mice living in room conditions (~ 2.5 years). However, we found that Sdh chKO mice survived longer than WT control mice in each of the 3 experimental groups (**Table 1**). When data from 3 groups were combined, the lifespan differences between Sdh chKO and WT mice were statistically significant (P<0.0001 by Log-rank (Mantel-Cox) test and P=0.0024 by Gehan-Breslow-Wilcoxon test. **Fig.1**). The rate of hypoxic death/moribund conditions in WT mice, estimated by Hazard ratio, was 11.39 (95% CI of 3.419 to 37.95) and 4.65 (95% CI of 1.606 to 13.47) fold higher than Sdh chKO mice by Mantel-Haenszel and logrank methods, respectively. Four WT mice and 1 Sdh bc mice were euthanized due to the development of morbid conditions that required euthanasia, as per IACUC protocols. The survival difference Sdh chKO and WT mice was statistically significant even when the mice euthanized for moribund conditions were excluded from the analysis (P=0.0037 Log-rank (Mantel-Cox) test and P=0.0304 Gehan-Breslow-Wilcoxon test).

**Figure 1.**
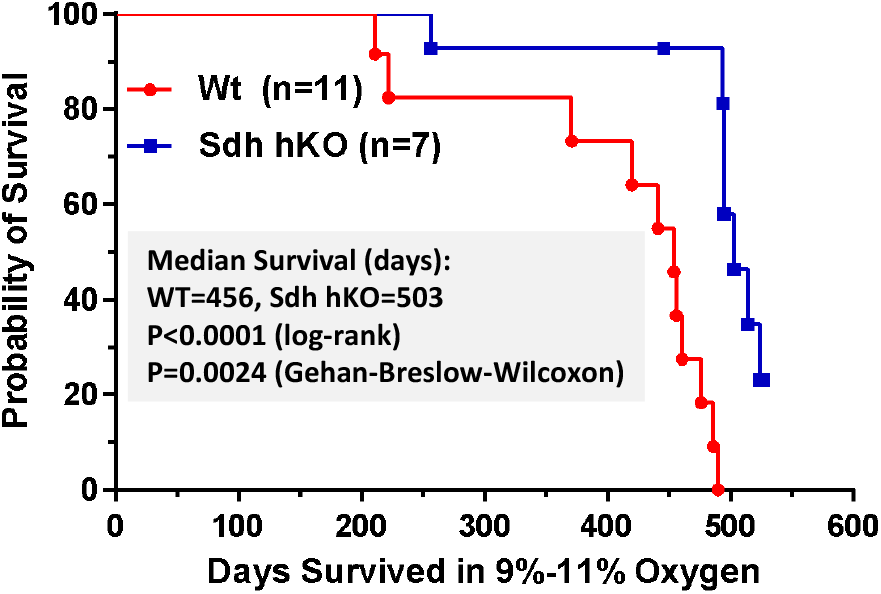
Kaplan-Meier survival curves of WT and Sdh chKO male mice from 3 groups (n=15 WT and n=15 chKO). The curve differences are statistically significant by Log-rank (Mantel-Cox) and Gehan-Breslow-Wilcoxon tests. Survival is measured by the total number of days until spontaneous death or the development of morbidities that require euthanasia (Table 1) in chronic hypoxia. Four WT and eight chKO mice are censored from the analysis since these mice did not complete the hypoxic survival end points (See Table 1).

Necropsy of mice revealed no specific causes to explain early death or development of moribund conditions, but showed congestion and enlarged spleen and heart which are expected under chronic hypoxia. Chronic hypoxia is associated with the development of pulmonary hypertension and right ventricular hypertrophy. We assessed right ventricular hypertrophy by Fulton index in experimental group 2, and found no statistically significant differences between Sdh and WT control mice (**Supp. Fig. 3**). This result suggests that pulmonary hypertension differences do not probably explain the differential survival between the Sdh chKO and WT mice.

### Sdh hKO mice show evidence of reduced RBC regeneration and lower red cell distribution width (RDW)

To examine whether erythrocyte numbers could explain the survival differences between the two genotypes, we analyzed CBC variables from 3 groups of Sdh hKO male mice and WT controls using 2way ANOVA test. No statistically significant differences were observed in erythrocyte numbers or hemoglobin (HGB) levels in any group (**Fig. 2A**). Reduced hematocrit (HCT) (**Fig. 2A**), mean corpuscle volume (MCV), mean corpuscular hemoglobin (MCH) (**Fig. 2B**), RDW-CV (**Fig. 2C**), reticulocyte percentage (Ret%) and immature reticulocyte fraction (IRF) (**Fig. 2D**) were also observed Sdh chKO mice in 1 of 3 groups (**Fig. 2**). The most statistically significant differences between the genotypes were observed in RDW-SD and 1SD-RDW which showed a reduction in Sdh chKO mice in 2 of 3 groups (**Fig. 2C**).

**Figure 2.**
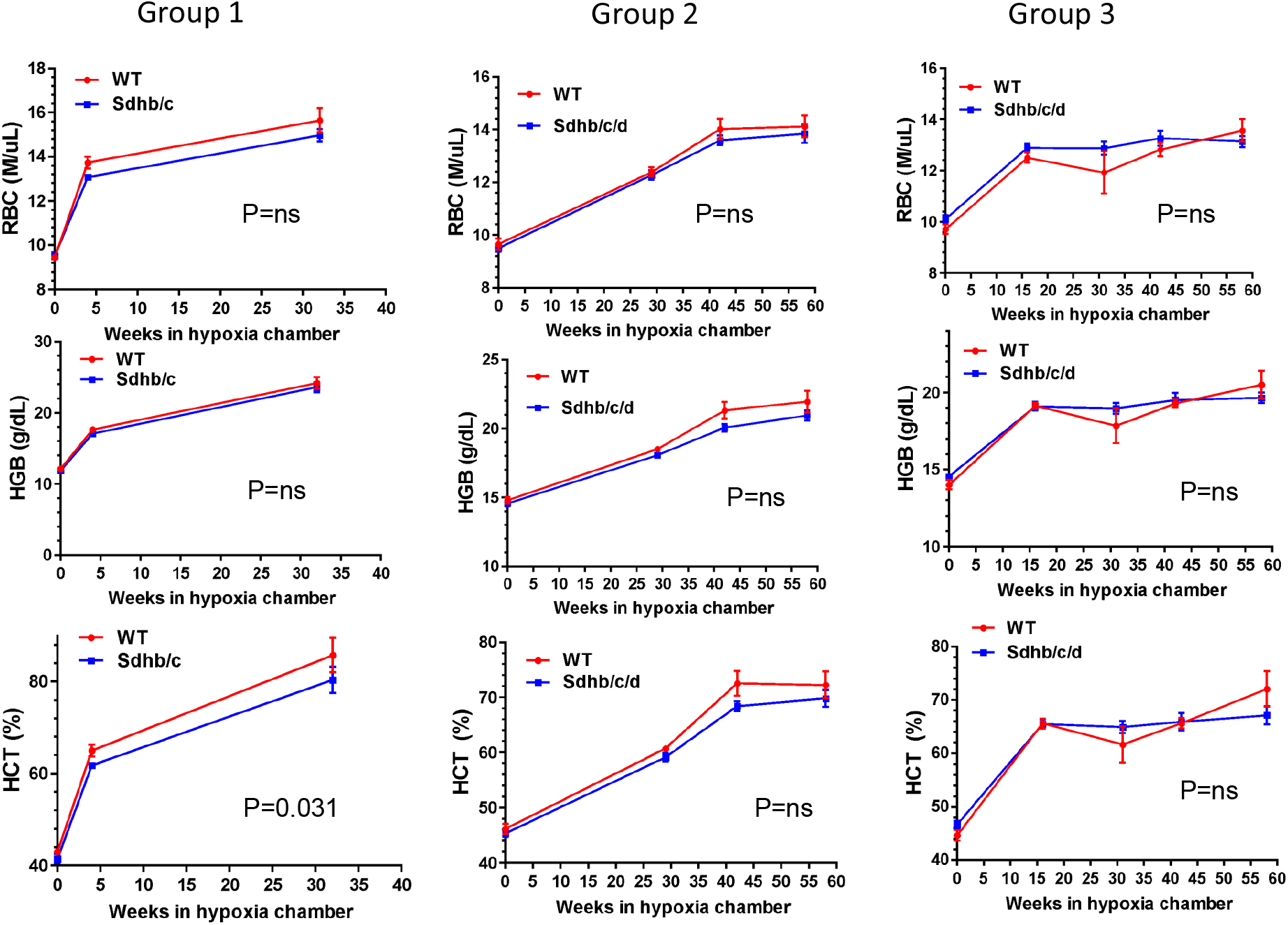
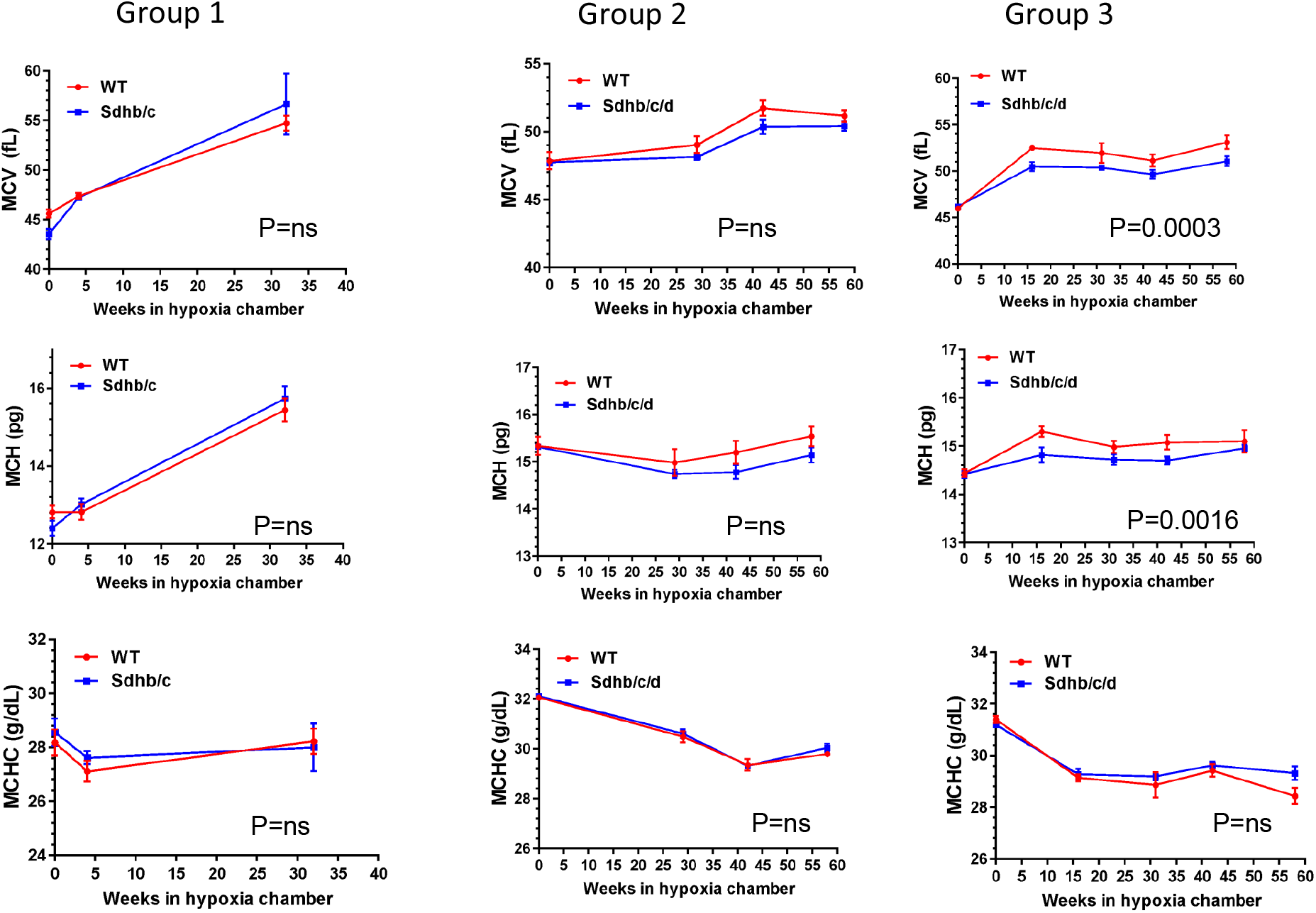
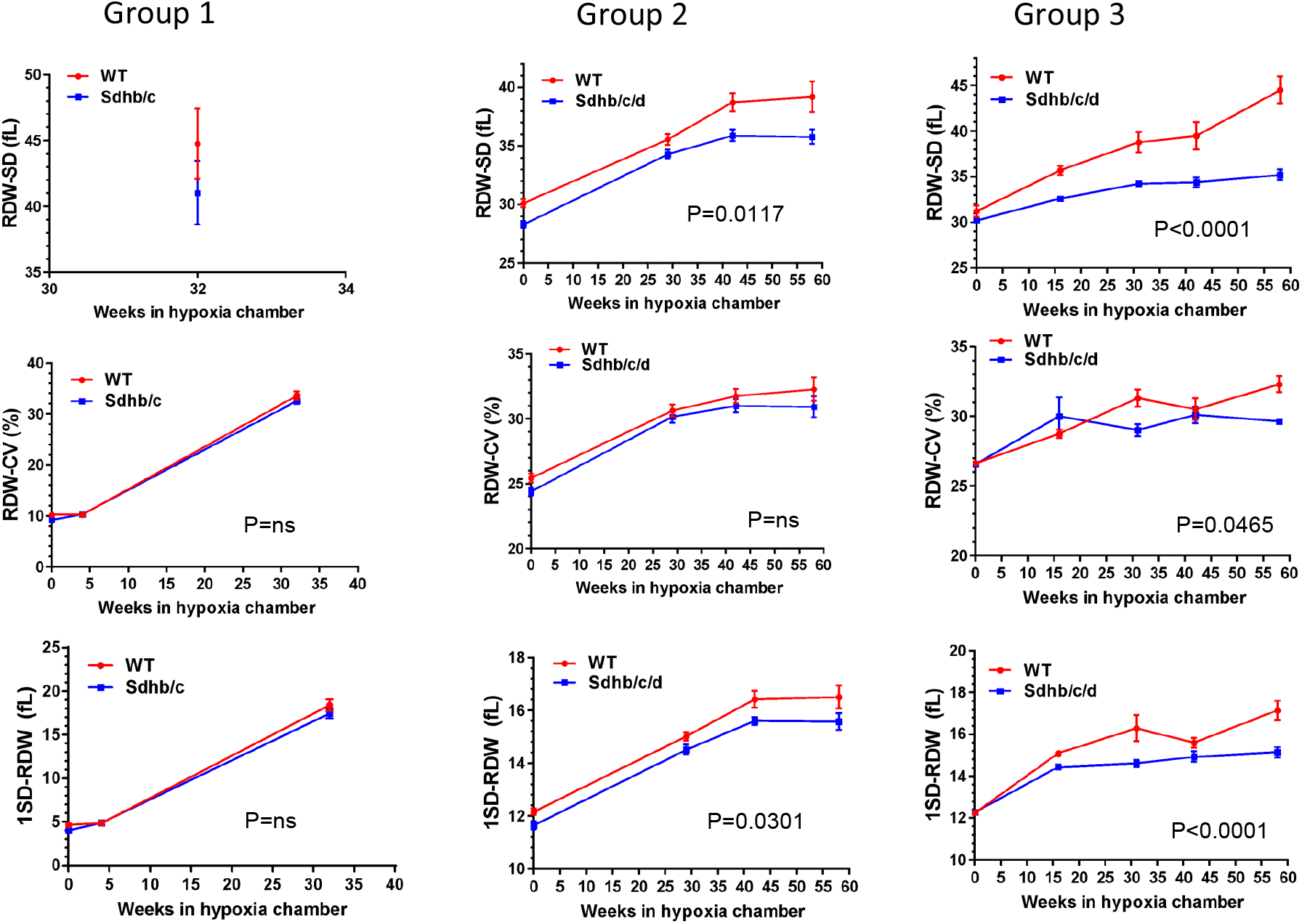
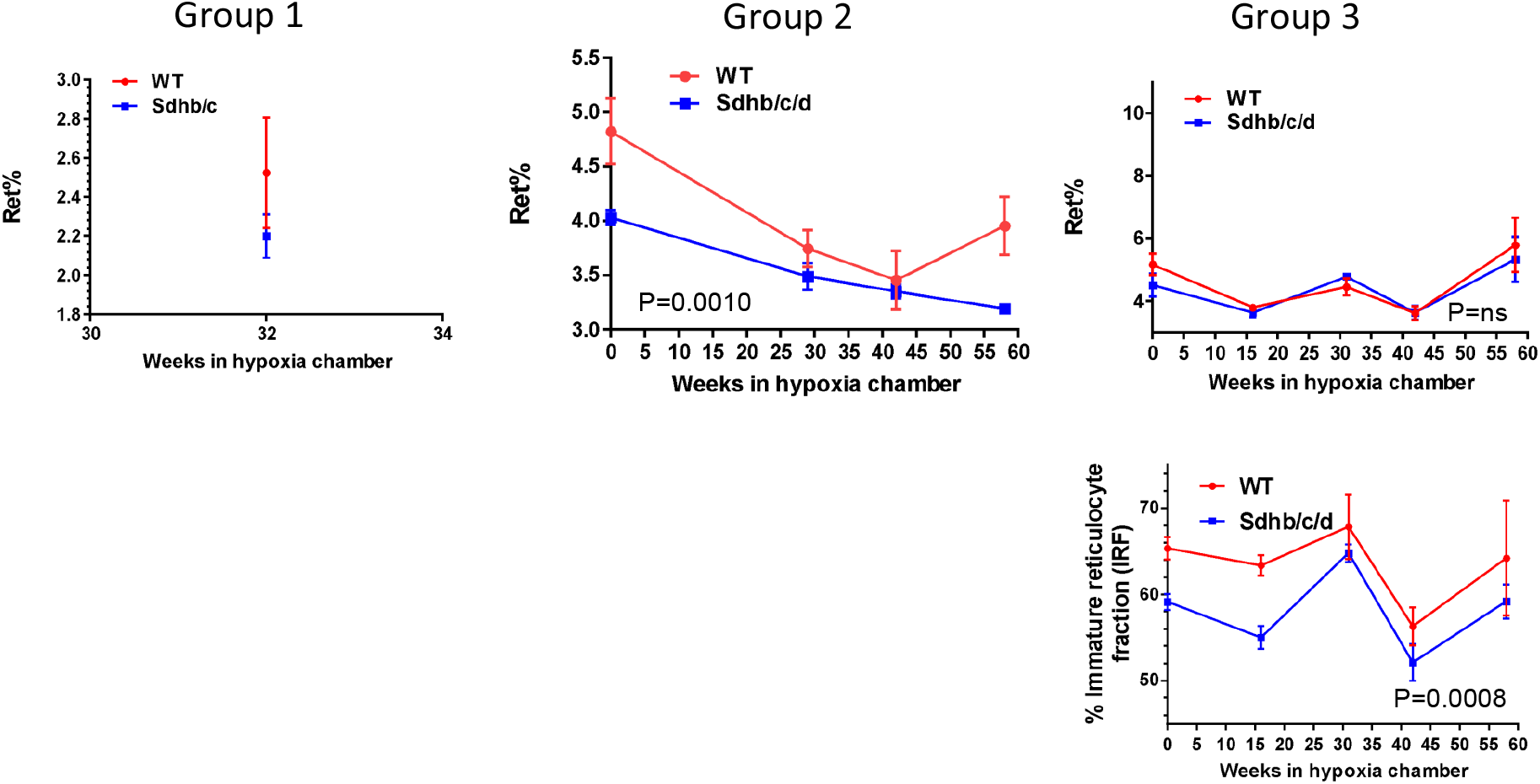
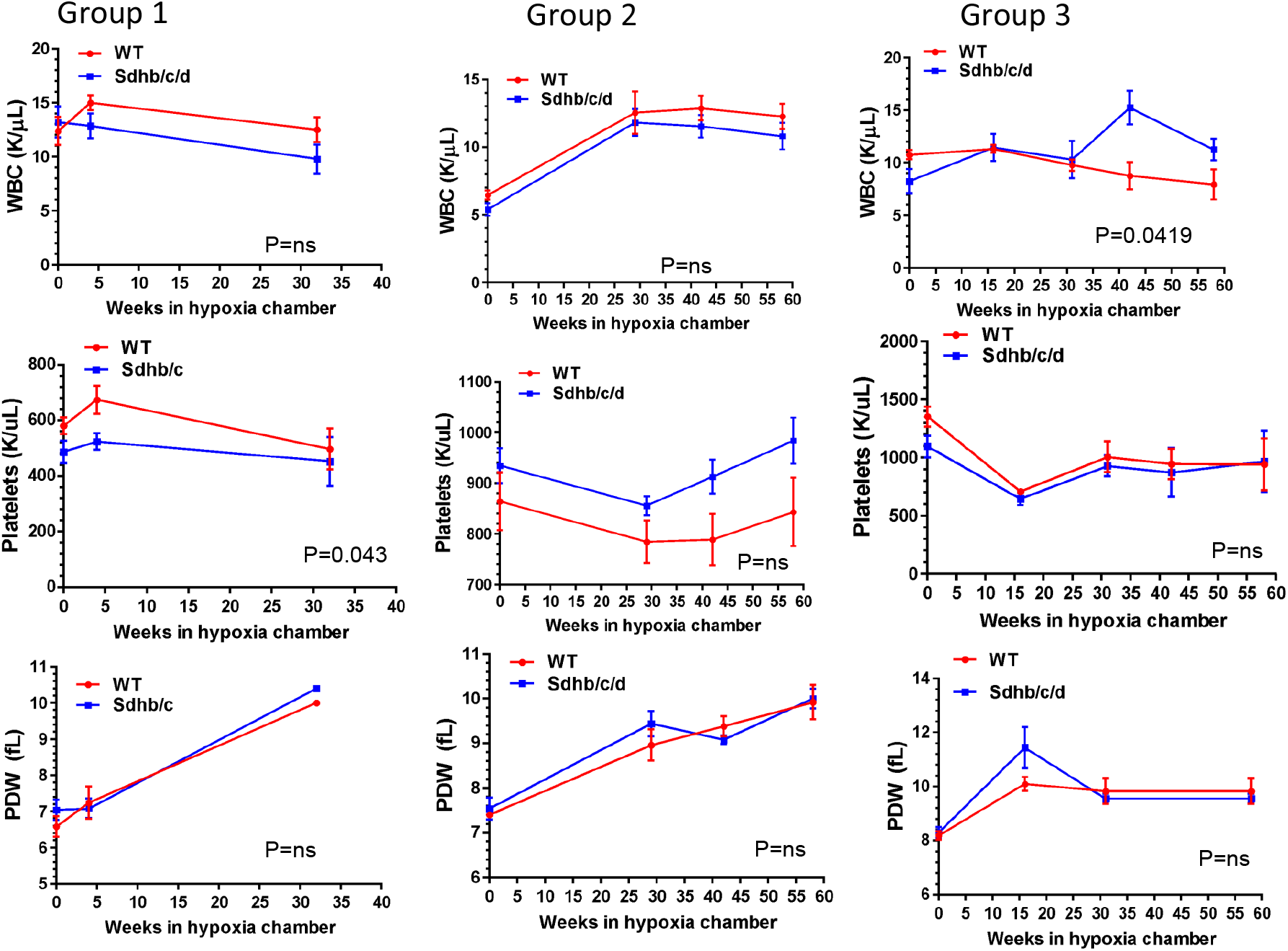
Erythrocyte parameters in Sdh chKO (Sdh bc in Group 1 and Sdh bcd in Groups 2 and 3) and WT control mice under chronic hypoxia. **(A)** RBC numbers, hemoglobin and hematocrit; **(B)** Mean corpuscle volume (MCV), mean corpuscle hemoglobin (MCH) and mean corpuscle hemoglobin concentration (MCHC); **(C)** Measures of RBC size variation including RDW-SD, RDW-CV and 1 standard deviation of RDW (1SD-RDW) and **(D)** Reticulocyte percentage (Ret%) and immature reticulocyte fraction (IRF). **(E)** White blood cell (WBC), platelet and platelet distribution width (PDW). Each time point contains 3-5 male mice. P values are calculated by 2way ANOVA using time and genotype as variables. Missing parameters in Group 1 (RDW-SD, Ret% and IRF) and Group 2 (IRF) were not reported in earlier CBC outputs. (HGB results of Groups 1 and 2 were originally shown in Sharma et al. (2017)[56], and included here for comprehensive analysis.)

Analysis of the combined data from all 3 groups showed the most statistically significant differences in RDW-SD, RDW-CV, 1SD-RDW and IRF both in normoxia and hypoxia with lower values observed in Sdh chKO mice relative to WT control mice (**Table 2**). Borderline statistically significant differences were seen in hematocrit and MCV in hypoxia and reticulocyte percentage in normoxia. No statistically significant differences were seen in the numbers of white blood cells, platelets and in platelet distribution width (PDW) between Sdh hKO and WT mice in normoxia or hypoxia (**Fig. 2E** and **Table 2**). Collectively, the differences in CBC parameters point to reduced erythropoietic activity and RBC regeneration by partial loss of Sdh, especially in hypoxia.

**Table 2.** Differences in RBC regeneration indices between Sdh chKO and WT male mice in baseline normoxia and/or chronic hypobaric hypoxia.

| Parameter | Normoxia (P values)<sup>#</sup> | Hypoxia (P values)<sup>#</sup> |
| --- | --- | --- |
| RBC (m/uL) | ns | ns |
| HGB (g/dL) | ns | ns |
| HCT (%) | ns | 0.0251 |
| MCV (fL) | ns | 0.0404 |
| MCH (pg) | ns | ns |
| MCHC (g/dL) | ns | ns |
| RDW-SD (fL) | 0.0023 | <0.0001<sup>a</sup> |
| RDW-CV (%) | 0.0027 | 0.0076 |
| 1SD-RDW (fL) | 0.0013 | <0.0001 |
| Ret% | 0.0257 | ns |
| Immature reticulocyte fraction (IRF) | 0.0159<sup>*,&</sup> | 0.0083<sup>*</sup> |
| WBC (K/uL) | ns | ns |
| Platelets (K/uL) | ns | ns |
| PDW (fL) | ns | ns |

## Discussion

In this study, we show that chKO mutations in Sdh genes prolong healthy survival by ~10% under chronic hypoxia and reduce multiple measures of RBC anisocytosis including RDW-SD, RDW-CV and 1SD-RDW. Other parameters related to RBC regeneration including IRF, HCT and MCV also show evidence of reductions in Sdh KO mice compared to WT. We find no evidence of PGL tumor development in mice even with chronic lifelong hypoxia exposure, in agreement with a recent study [58]. These findings collectively show a previously unrecognized role for Sdh in regulation of erythroid regeneration both in normoxia and hypoxia. To our knowledge, this is also the first mammalian study showing a hypoxia-survival benefit upon partial constitutional loss of Sdh, suggesting a role for Sdh in cellular and organismal adaptation to hypoxia in mice.

Detailed studies in *Ascaris Suum*, a helminthic parasite, show that Sdh is active in spore forms which respire atmospheric O_2_, but inactive in the adult forms which live in the hypoxic environment of the host intestine. The adult parasite instead uses fumarate reductase (Frd), which catalyzes the reverse reaction of Sdh [59]. A hypoxic switch in Sdh genes also controls respiration in *Mycobacterium Tuberculosis* [60]. ATP producing eukaryotic mitochondria use Frd rather than Sdh under limited O_2_ conditions [61]. Flies resistant to hypoxia have reduced complex II activity levels compared to the control flies [62]. Anoxic environments (N2 or CO_2_) lead to decreased transcript expression of the 3 of four SDH subunit genes, by promoter methylation in maize [63]. Our findings combined with these studies suggest that inhibition of Sdh is a universal theme in organismal adaptation to hypoxia/anoxia across diverse organisms including mammals.

Identification of the molecular mechanisms linking reduced Sdh to organismal tolerance to hypoxia requires further studies. There is already evidence that loss of SDH triggers hypoxia adaptation pathways in human PGL tumors. The *SDHD* gene is subject to maternal imprinting (inactivation) in hypoxia-sensitive carotid body chief cells, because only a paternal transmission, but not maternal transmission, of the mutated *SDHD* gene predisposes to paraganglioma tumors. This finding raises the hypothesis that partial loss of SDH activity by genomic imprinting is physiologically employed to facilitate hypoxia sensing and/or adaptation in carotid body cells [64]. We recently showed that inhibition of complex II by atpenin A5 triggers hypoxic gene expression and RNA editing by APOBEC3A and APOBEC3G cytidine deaminases independently of HIF1 in monocytes and natural killer (NK) cells, respectively [56, 65]. The *SDHB* and *SDHA* genes acquire nonsense/missense RNA editing by APOBEC3A in monocytes subjected to cellular crowding and hypoxia [66, 67]. RNA editing by APOBEC3G in NK cells is induced by cellular crowding and hypoxia and promotes Warburg-like metabolic remodeling by suppressing O_2_ consumption relative to glycolysis [65]. Perhaps, the inhibition of Sdh in mice activates similar HIF-independent hypoxia-adaptation pathways including gene expression, and RNA editing, and possibly other adaptive pathways that remain to be discovered.

Importantly, our findings show a mitochondrial basis for the association between high RDW and mortality. Previous research has established that inhibition of mitochondrial respiration antagonizes the hypoxic stabilization of HIF-α [68, 69], the key molecular event driving the synthesis of erythropoietin that stimulates RBC regeneration in bone marrow. Pharmacologic inhibition of complex II by atpenin A5 reduces the stabilization of HIF-α in cancer cell lines in hypoxia, and reduces baseline O_2_ consumption [56, 70]. Atpenin A5 is a highly potent complex II inhibitor of ubiquinone binding that occurs at the interface of Sdhb, Sdhc and Sdhd subunits [71, 72]. Therefore, we suggest that partial loss of Sdh in hKO mice reduces mitochondrial O_2_ consumption and dampens HIF-mediated erythropoietic activity, leading to reduced RDW in hypoxia.

Our study has certain limitations including the lack of a specific heart or lung disease in which high RDW has been associated with early mortality in clinical studies, and indeterminate cause of death in hypoxic mice. Also, these findings remain to be extended to female mice. Further studies are required to close these knowledge gaps in the future.

In summary, our findings provide evidence that Sdh plays a role in regenerative erythrocyte anisocytosis and organismal survival in mice under chronic hypoxia. Our data support a model that upon systemic hypoxia, continued mitochondrial O_2_ consumption leads to cellular O_2_ deprivation, increased erythropoiesis and high RDW. This unchecked O_2_ consumption ultimately exhausts intracellular O_2_ to cause cellular injury, organ failure and death. Suppressing Sdh reduces O_2_ consumption, mitigates cellular hypoxia, blunts RDW increase and triggers HIF-independent hypoxia adaptation pathways to promote organismal tolerance to chronic hypoxia **(Fig.3)**. We hypothesize that high RDW is merely a surrogate biomarker for organismal hypoxia which is regulated by mitochondrial O_2_ consumption and is the ultimate driver of cell death, organ failure and mortality. Therefore, therapeutic targeting of SDH may be beneficial to reduce high RDW-associated mortality in hypoxic diseases by enhancing systemic adaptation to hypoxia.

**Figure 3.**
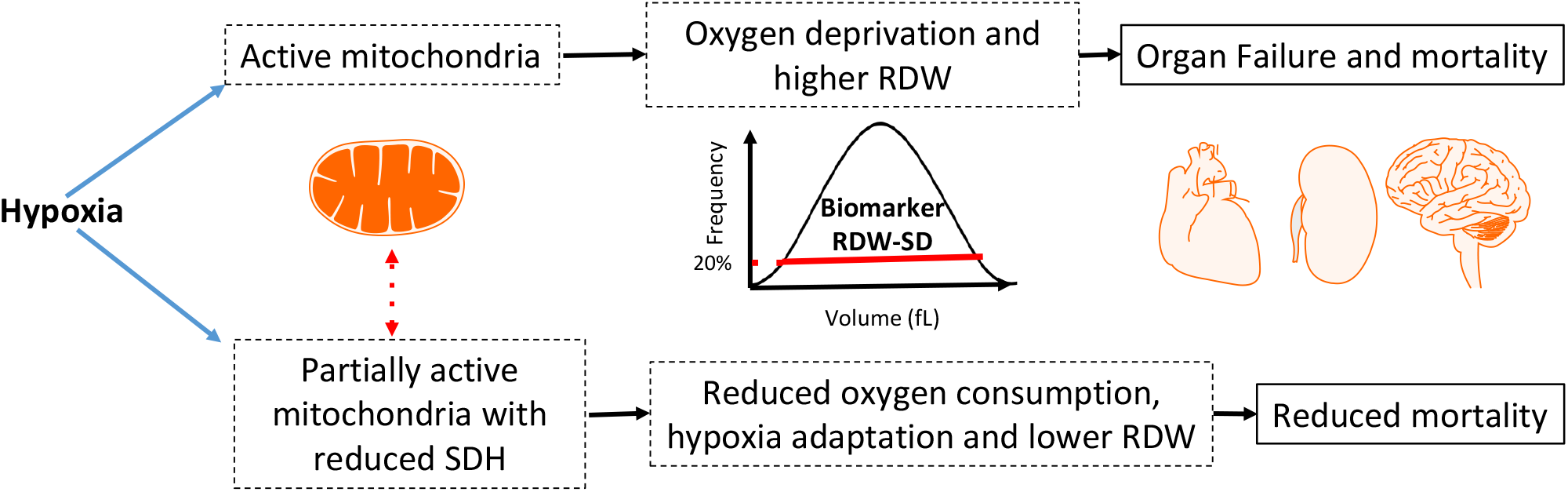
A mitochondrial basis for the association between high RDW and mortality. When oxygen is limited, fully active mitochondria exhausts the remaining oxygen leading to (a) erythrocyte regeneration and high RDW and (b) reduced cellular viability, organ failure and mortality. Inhibition of SDH reduces oxygen consumption and RDW levels, and triggers cellular hypoxia adaptation pathways leading to improved survival.

## Supplementary Figure Legends

**Supp. Fig. 1.** MRI analysis of hypoxia exposed WT (**A**) and BC chKO (**B**) show non-specific findings but no definite evidence of tumor development.

**Supp. Fig. 2.** Microscopic examination of lungs (**A**) and adrenal glands (**B**) show no morphologic evidence of vascular abnormalities or pheochromocytoma. Arrows show lung vessels without morphologic evidence of intimal hyperplasia in lung (**A**) or point to adrenal medulla (**B**), surrounded by adrenal cortex in adrenal gland.

**Supp. Fig. 3.** Fulton index (ratio of weight of right ventricle to sum of left ventricle and septum) in group 2 mice showed no statistically significant difference.

## Acknowledgement

This research was supported by startup funds from the Departments of Pathology (BEB), and and by National Cancer Institute (NCI) Grant (P30CA016056) involving the use of RPCCC’s Laboratory Animal Shared Resource (LASR) and Gene Targeting & Transgenic Shared Resource (GeTT).

## Contributions

BEB designed the study with contributions from DT and SS. DT performed mice handling, care, identification and breeding. DT, SS and LC performed mice physical evaluations, blood draws and necropsy. MS performed MRI analysis. BEB performed the stastitical analysis, prepared the figures and wrote the manuscript. All authors received the manuscript and agreed on the authorship.

## References

1. Constantino BT: Red cell distribution width, revisited. Laboratory Medicine 2013, 44(2):e2–e9.

2. Lippi G, Salvagno GL, Guidi GC: Red Blood Cell Distribution Width Is Significantly Associated With Aging and Gender. Clinical chemistry and laboratory medicine 2014, 52(9):e197–e199.

3. Patel KV, Ferrucci L, Ershler WB, Longo DL, Guralnik JM: Red blood cell distribution width and the risk of death in middle-aged and older adults. Archives of internal medicine 2009, 169(5):515–523.

4. Perlstein TS, Weuve J, Pfeffer MA, Beckman JA: Red blood cell distribution width and mortality risk in a community-based prospective cohort. Archives of Internal Medicine 2009, 169(6):588–594.

5. Felker GM, Allen LA, Pocock SJ, Shaw LK, McMurray JJV, Pfeffer MA, Swedberg K, Wang D, Yusuf S, Michelson EL: Red cell distribution width as a novel prognostic marker in heart failure: data from the CHARM Program and the Duke Databank. Journal of the American College of Cardiology 2007, 50(1):40–47.

6. Allen LA, Felker GM, Mehra MR, Chiong JR, Dunlap SH, Ghali JK, Lenihan DJ, Oren RM, Wagoner LE, Schwartz TA: Validation and potential mechanisms of red cell distribution width as a prognostic marker in heart failure. Journal of cardiac failure 2010, 16(3):230–238.

7. Dabbah S, Hammerman H, Markiewicz W, Aronson D: Relation between red cell distribution width and clinical outcomes after acute myocardial infarction. The American journal of cardiology 2010, 105(3):312–317.

8. Ye Z, Smith C, Kullo IJ: Usefulness of red cell distribution width to predict mortality in patients with peripheral artery disease. The American journal of cardiology 2011, 107(8):1241–1245.

9. Montagnana M, Danese E: Red cell distribution width and cancer. Annals of translational medicine 2016, 4(20).

10. Hampole CV, Mehrotra AK, Thenappan T, Gomberg-Maitland M, Shah SJ: Usefulness of red cell distribution width as a prognostic marker in pulmonary hypertension. The American journal of cardiology 2009, 104(6):868–872.

11. Zorlu A, Bektasoglu G, Guven FMK, Dogan OT, Gucuk E, Ege MR, Altay H, Cinar Z, Tandogan I, Yilmaz MB: Usefulness of admission red cell distribution width as a predictor of early mortality in patients with acute pulmonary embolism. The American journal of cardiology 2012, 109(1):128–134.

12. Lee JH, Chung HJ, Kim K, Jo YH, Rhee JE, Kim YJ, Kang KW: Red cell distribution width as a prognostic marker in patients with community-acquired pneumonia. The American journal of emergency medicine 2013, 31(1):72–79.

13. Braun E, Domany E, Kenig Y, Mazor Y, Makhoul BF, Azzam ZS: Elevated red cell distribution width predicts poor outcome in young patients with community acquired pneumonia. Critical care 2011, 15(4):1–9.

14. Foy BH, Carlson JC, Reinertsen E, Valls RPI, Lopez RP, Palanques-Tost E, Mow C, Westover MB, Aguirre AD, Higgins JM: Association of red blood cell distribution width with mortality risk in hospitalized adults with SARS-CoV-2 infection. JAMA Network Open 2020, 3(9):e2022058–e2022058.

15. Seyhan EC, Özgül MA, Tutar N, Ömür Im, Uysal A, Altın S: Red blood cell distribution and survival in patients with chronic obstructive pulmonary disease. COPD: Journal of Chronic Obstructive Pulmonary Disease 2013, 10(4):416–424.

16. Epstein D, Nasser R, Mashiach T, Azzam ZS, Berger G: Increased red cell distribution width: A novel predictor of adverse outcome in patients hospitalized due to acute exacerbation of chronic obstructive pulmonary disease. Respiratory medicine 2018, 136:1–7.

17. Wang B, Gong Y, Ying B, Cheng B: Relation between red cell distribution width and mortality in critically ill patients with acute respiratory distress syndrome. BioMed research international 2019, 2019.

18. Kim J, Kim YD, Song T-J, Park JH, Lee HS, Nam CM, Nam HS, Heo JH: Red blood cell distribution width is associated with poor clinical outcome in acute cerebral infarction. Thrombosis and haemostasis 2012, 108(08):349–356.

19. Ani C, Ovbiagele B: Elevated red blood cell distribution width predicts mortality in persons with known stroke. Journal of the neurological sciences 2009, 277(1-2):103–108.

20. Wang F, Pan W, Pan S, Ge J, Wang S, Chen M: Red cell distribution width as a novel predictor of mortality in ICU patients. Annals of medicine 2011, 43(1):40–46.

21. Bazick HS, Chang D, Mahadevappa K, Gibbons FK, Christopher KB: Red Cell Distribution Width and all cause mortality in critically ill patients. Critical care medicine 2011, 39(8):1913.

22. Majercik S, Fox J, Knight S, Horne BD: Red cell distribution width is predictive of mortality in trauma patients. Journal of Trauma and Acute Care Surgery 2013, 74(4):1021–1026.

23. Garbharran U, Chinthapalli S, Hopper I, George M, Back DL, Dockery F: Red cell distribution width is an independent predictor of mortality in hip fracture. Age and ageing 2013, 42(2):258–261.

24. Jo YH, Kim K, Lee JH, Kang C, Kim T, Park H-M, Kang KW, Kim J, Rhee JE: Red cell distribution width is a prognostic factor in severe sepsis and septic shock. The American journal of emergency medicine 2013, 31(3):545–548.

25. Ku NS, Kim H-w, Oh HJ, Kim YC, Kim MH, Song JE, Oh DH, Ahn JY, Kim SB, Jeong SJ: Red blood cell distribution width is an independent predictor of mortality in patients with gram-negative bacteremia. Shock 2012, 38(2):123–127.

26. Şenol K, Saylam B, Kocaay F, Tez M: Red cell distribution width as a predictor of mortality in acute pancreatitis. The American journal of emergency medicine 2013, 31(4):687–689.

27. Vashistha T, Streja E, Molnar MZ, Rhee CM, Moradi H, Soohoo M, Kovesdy CP, Kalantar-Zadeh K: Red cell distribution width and mortality in hemodialysis patients. American journal of kidney diseases 2016, 68(1):110–121.

28. Mucsi I, Ujszaszi A, Czira ME, Novak M, Molnar MZ: Red cell distribution width is associated with mortality in kidney transplant recipients. International urology and nephrology 2014, 46(3):641–651.

29. Cavusoglu E, Chopra V, Gupta A, Battala VR, Poludasu S, Eng C, Marmur JD: Relation between red blood cell distribution width (RDW) and all-cause mortality at two years in an unselected population referred for coronary angiography. International journal of cardiology 2010, 141(2):141–146.

30. Uyarel H, Ergelen M, Cicek G, Kaya MG, Ayhan E, Turkkan C, Yildirim E, Kirbas V, Onturk ET, Erer HB: Red cell distribution width as a novel prognostic marker in patients undergoing primary angioplasty for acute myocardial infarction. Coronary artery disease 2011, 22(3): 138-144.

31. Lam AP, Gundabolu K, Sridharan A, Jain R, Msaouel P, Chrysofakis G, Yu Y, Friedman E, Price E, Schrier S: Multiplicative interaction between mean corpuscular volume and red cell distribution width in predicting mortality of elderly patients with and without anemia. American journal of hematology 2013, 88(11):E245–E249.

32. Núñez J, Núñez E, Rizopoulos D, Miñana G, Bodí V, Bondanza L, Husser O, Merlos P, Santas E, Pascual-Figal D: Red blood cell distribution width is longitudinally associated with mortality and anemia in heart failure patients. Circulation Journal 2014, 78(2):410–418.

33. Lv H, Zhang L, Long A, Mao Z, Shen J, Yin P, Li M, Zeng C, Zhang L, Tang P: Red cell distribution width as an independent predictor of long-term mortality in hip fracture patients: a prospective cohort study. Journal of Bone and Mineral Research 2016, 31(1):223–233.

34. Shah N, Pahuja M, Pant S, Handa A, Agarwal V, Patel N, Dusaj R: Red cell distribution width and risk of cardiovascular mortality: Insights from National Health and Nutrition Examination Survey (NHANES)-III. International journal of cardiology 2017, 232:105–110.

35. Patel KV, Semba RD, Ferrucci L, Newman AB, Fried LP, Wallace RB, Bandinelli S, Phillips CS, Yu B, Connelly S: Red cell distribution width and mortality in older adults: a meta-analysis. Journals of Gerontology Series A: Biomedical Sciences and Medical Sciences 2009, 65(3):258–265.

36. Lippi G, Targher G, Montagnana M, Salvagno GL, Zoppini G, Guidi GC: Relation between red blood cell distribution width and inflammatory biomarkers in a large cohort of unselected outpatients. Archives of Pathology & Laboratory Medicine 2009, 133(4):628–632.

37. Emans ME, Gaillard CA, Pfister R, Tanck MW, Boekholdt SM, Wareham NJ, Khaw K-T: Red cell distribution width is associated with physical inactivity and heart failure, independent of established risk factors, inflammation or iron metabolism; the EPIC—Norfolk study. International journal of cardiology 2013, 168(4):3550–3555.

38. Lappé JM, Horne BD, Shah SH, May HT, Muhlestein JB, Lappé DL, Kfoury AG, Carlquist JF, Budge D, Alharethi R: Red cell distribution width, C-reactive protein, the complete blood count, and mortality in patients with coronary disease and a normal comparison population. Clinica chimica acta 2011, 412(23-24):2094–2099.

39. Patel KV, Mohanty JG, Kanapuru B, Hesdorffer C, Ershler WB, Rifkind JM: Association of the red cell distribution width with red blood cell deformability. In: Oxygen Transport to Tissue XXXIV. edn.: Springer; 2013: 211–216.

40. Vayá A, Rivera L, de la Espriella R, Sanchez F, Suescun M, Hernandez JL, Fácila L: Red blood cell distribution width and erythrocyte deformability in patients with acute myocardial infarction. Clinical hemorheology and microcirculation 2015, 59(2):107–114.

41. Sun H, Weaver CM: Decreased Iron Intake Parallels Rising Iron Deficiency Anemia and Related Mortality Rates in the US Population. The Journal of Nutrition 2021.

42. Yčas JW, Horrow JC, Horne BD: Persistent increase in red cell size distribution width after acute diseases: A biomarker of hypoxemia? Clinica Chimica Acta 2015, 448:107–117.

43. Tertemiz KC, Alpaydin AO, Sevinc C, Ellidokuz H, Acara AC, Cimrin A: Could “red cell distribution width” predict COPD severity? Revista Portuguesa de Pneumologia (English Edition) 2016, 22(4):196–201.

44. Karampitsakos T, Dimakou K, Papaioannou O, Chrysikos S, Kaponi M, Bouros D, Tzouvelekis A, Hillas G: The role of increased red cell distribution width as a negative prognostic marker in patients with COPD. Pulmonary pharmacology & therapeutics 2020, 60:101877.

45. Grant BJ, Kudalkar DP, Muti P, McCann SE, Trevisan M, Freudenheim JL, Schu HJ: Relation between lung function and RBC distribution width in a population-based study. Chest 2003, 124(2):494–500.

46. Thayer TE, Huang S, Levinson RT, Farber-Eger E, Assad TR, Huston JH, Mosley JD, Wells QS, Brittain EL: Unbiased Phenome-Wide Association Studies of Red Cell Distribution Width Identifies Key Associations with Pulmonary Hypertension. Ann Am Thorac Soc 2019, 16(5):589–598.

47. Xie J, Covassin N, Fan Z, Singh P, Gao W, Li G, Kara T, Somers VK: Association between hypoxemia and mortality in patients with COVID-19. Mayo Clinic Proceedings 2020, 95(6):1138–1147.

48. Franke K, Gassmann M, Wielockx B: Erythrocytosis: the HIF pathway in control. Blood, The Journal of the American Society of Hematology 2013, 122(7):1122–1128.

49. Favier J, Amar L, Gimenez-Roqueplo A-P: Paraganglioma and phaeochromocytoma: from genetics to personalized medicine. Nature Reviews Endocrinology 2015, 11(2):101–111.

50. Baysal BE, Ferrell RE, Willett-Brozick JE, Lawrence EC, Myssiorek D, Bosch A, van der Mey A, Taschner PE, Rubinstein WS, Myers EN et al: Mutations in SDHD, a mitochondrial complex II gene, in hereditary paraganglioma. Science (New York, NY) 2000, 287(5454):848–851.

51. Rodriguez-Cuevas H, Lau I, Rodriguez HP: High-altitude paragangliomas diagnostic and therapeutic considerations. Cancer 1986, 57(3):672–676.

52. Astrom K, Cohen JE, Willett-Brozick JE, Aston CE, Baysal BE: Altitude is a phenotypic modifier in hereditary paraganglioma type 1: evidence for an oxygen-sensing defect. Human genetics 2003, 113(3):228–237.

53. Cerecer-Gil NY, Figuera LE, Llamas FJ, Lara M, Escamilla JG, Ramos R, Estrada G, Hussain AK, Gaal J, Korpershoek E: Mutation of SDHB is a cause of hypoxia-related high-altitude paraganglioma. Clinical Cancer Research 2010, 16(16):4148–4154.

54. Castro-Vega LJ, Letouzé E, Burnichon N, Buffet A, Disderot P-H, Khalifa E, Loriot C, Elarouci N, Morin A, Menara M: Multi-omics analysis defines core genomic alterations in pheochromocytomas and paragangliomas. Nature communications 2015, 6(1):1–9.

55. Piruat JI, Millán-Uclés Á: Genetically modeled mice with mutations in mitochondrial metabolic enzymes for the study of cancer. Frontiers in oncology 2014, 4:200.

56. Sharma S, Wang J, Cortes Gomez E, Taggart RT, Baysal BE: Mitochondrial complex II regulates a distinct oxygen sensing mechanism in monocytes. Human molecular genetics 2017, 26(7):1328–1339.

57. Piruat JI, Pintado CO, Ortega-Sáenz P, Roche M, López-Barneo J: The mitochondrial SDHD gene is required for early embryogenesis, and its partial deficiency results in persistent carotid body glomus cell activation with full responsiveness to hypoxia. Molecular and cellular biology 2004, 24(24):10933–10940.

58. Al Khazal F, Kang S, Nelson Holte M, Choi DS, Singh R, Ortega-Sáenz P, López-Barneo J, Maher III LJ: Unexpected obesity, rather than tumorigenesis, in a conditional mouse model of mitochondrial complex II deficiency. The FASEB Journal 2021, 35(2):e21227.

59. Kita K, Hirawake H, Miyadera H, Amino H, Takeo S: Role of complex II in anaerobic respiration of the parasite mitochondria from Ascaris suum and Plasmodium falciparum. Biochimica et Biophysica Acta (BBA)-Bioenergetics 2002, 1553(1-2):123–139.

60. Hartman T, Weinrick B, Vilchèze C, Berney M, Tufariello J, Cook GM, Jacobs Jr WR: Succinate dehydrogenase is the regulator of respiration in Mycobacterium tuberculosis. PLoS Pathog 2014, 10(11):e1004510.

61. Müller M, Mentel M, van Hellemond JJ, Henze K, Woehle C, Gould SB, Yu R-Y, van der Giezen M, Tielens AG, Martin WF: Biochemistry and evolution of anaerobic energy metabolism in eukaryotes. Microbiology and Molecular Biology Reviews 2012, 76(2):444–495.

62. Ali SS, Hsiao M, Zhao HW, Dugan LL, Haddad GG, Zhou D: Hypoxia-adaptation involves mitochondrial metabolic depression and decreased ROS leakage. PloS one 2012, 7(5):e36801.

63. Eprintsev AT, Fedorin DN, Dobychina MA, Igamberdiev AU: Expression and promoter methylation of succinate dehydrogenase and fumarase genes in maize under anoxic conditions. Journal of plant physiology 2017, 216:197–201.

64. Baysal BE: Genomic imprinting and environment in hereditary paraganglioma. American journal of medical geneticsPart C, Seminars in medical genetics 2004, 129C(1):85–90.

65. Sharma S, Wang J, Alqassim E, Portwood S, Cortes Gomez E, Maguire O, Basse PH, Wang ES, Segal BH, Baysal BE: Mitochondrial hypoxic stress induces widespread RNA editing by APOBEC3G in natural killer cells. Genome biology 2019, 20(1):37-019-1651-1651.

66. Sharma S, Patnaik SK, Taggart RT, Kannisto ED, Enriquez SM, Gollnick P, Baysal BE: APOBEC3A cytidine deaminase induces RNA editing in monocytes and macrophages. Nature communications 2015, 6(1):1–15.

67. Baysal BE, De Jong K, Liu B, Wang J, Patnaik SK, Wallace PK, Taggart RT: Hypoxia-inducible C-to-U coding RNA editing downregulates SDHB in monocytes. PeerJ 2013, 1:e152.

68. Lin X, David CA, Donnelly JB, Michaelides M, Chandel NS, Huang X, Warrior U, Weinberg F, Tormos KV, Fesik SW: A chemical genomics screen highlights the essential role of mitochondria in HIF-1 regulation. Proceedings of the National Academy of Sciences 2008, 105(1):174–179.

69. Taylor CT: Mitochondria and cellular oxygen sensing in the HIF pathway. Biochemical journal 2008, 409(1):19–26.

70. Quinlan CL, Orr AL, Perevoshchikova IV, Treberg JR, Ackrell BA, Brand MD: Mitochondrial complex II can generate reactive oxygen species at high rates in both the forward and reverse reactions. Journal of Biological Chemistry 2012, 287(32):27255–27264.

71. Miyadera H, Shiomi K, Ui H, Yamaguchi Y, Masuma R, Tomoda H, Miyoshi H, Osanai A, Kita K, Ōmura S: Atpenins, potent and specific inhibitors of mitochondrial complex II (succinate-ubiquinone oxidoreductase). Proceedings of the National Academy of Sciences 2003, 100(2):473–477.

72. Wang H, Huwaimel B, Verma K, Miller J, Germain TM, Kinarivala N, Pappas D, Brookes PS, Trippier PC: Synthesis and antineoplastic evaluation of mitochondrial complex II (succinate dehydrogenase) inhibitors derived from atpenin A5. ChemMedChem 2017, 12(13):1033–1044.

