## Supplementary Figures 1-3 for "Partial loss of succinate dehydrogenase reduces high red cell distribution width and promotes healthy survival in chronically hypoxic mice"

# 197 WT

Supp. Fig.1A

Superior | Inferior 1 | Inferior 2

Coronal

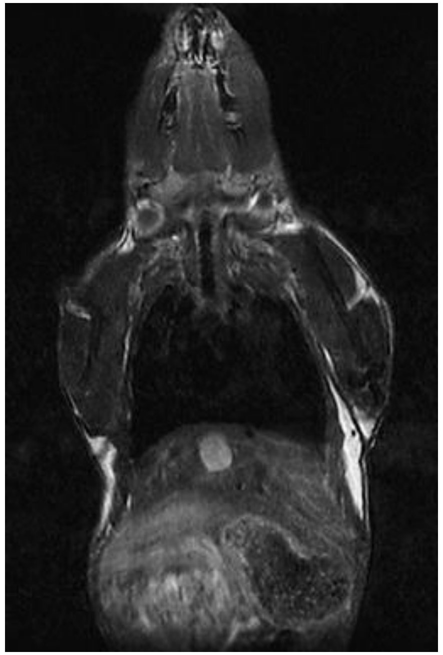

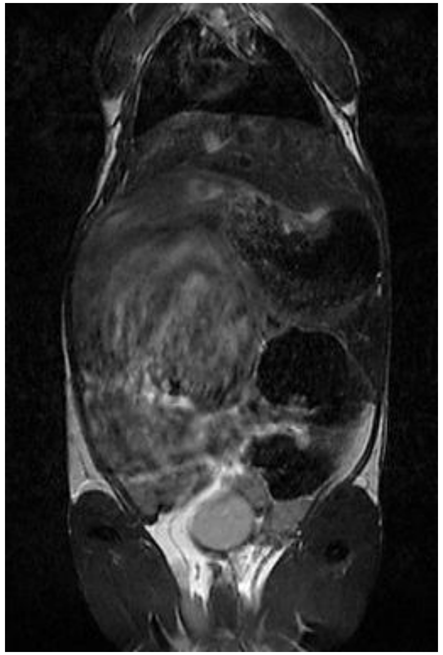

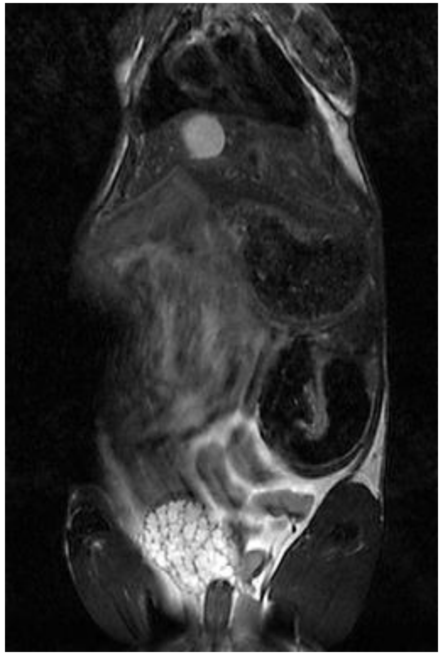

No unusual features noticed in superior segment.

Axial

Enlarged seminal vesicles

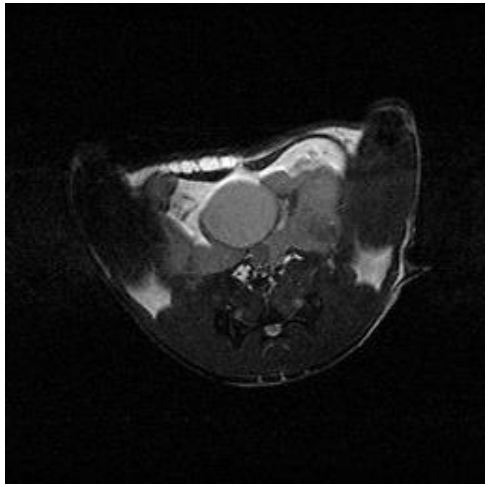

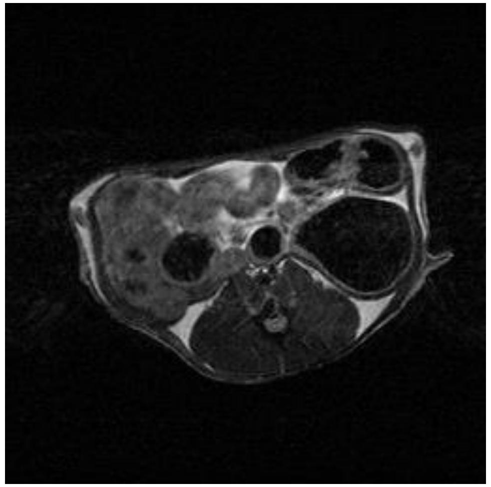

# 145 BC

Supp. Fig.1B

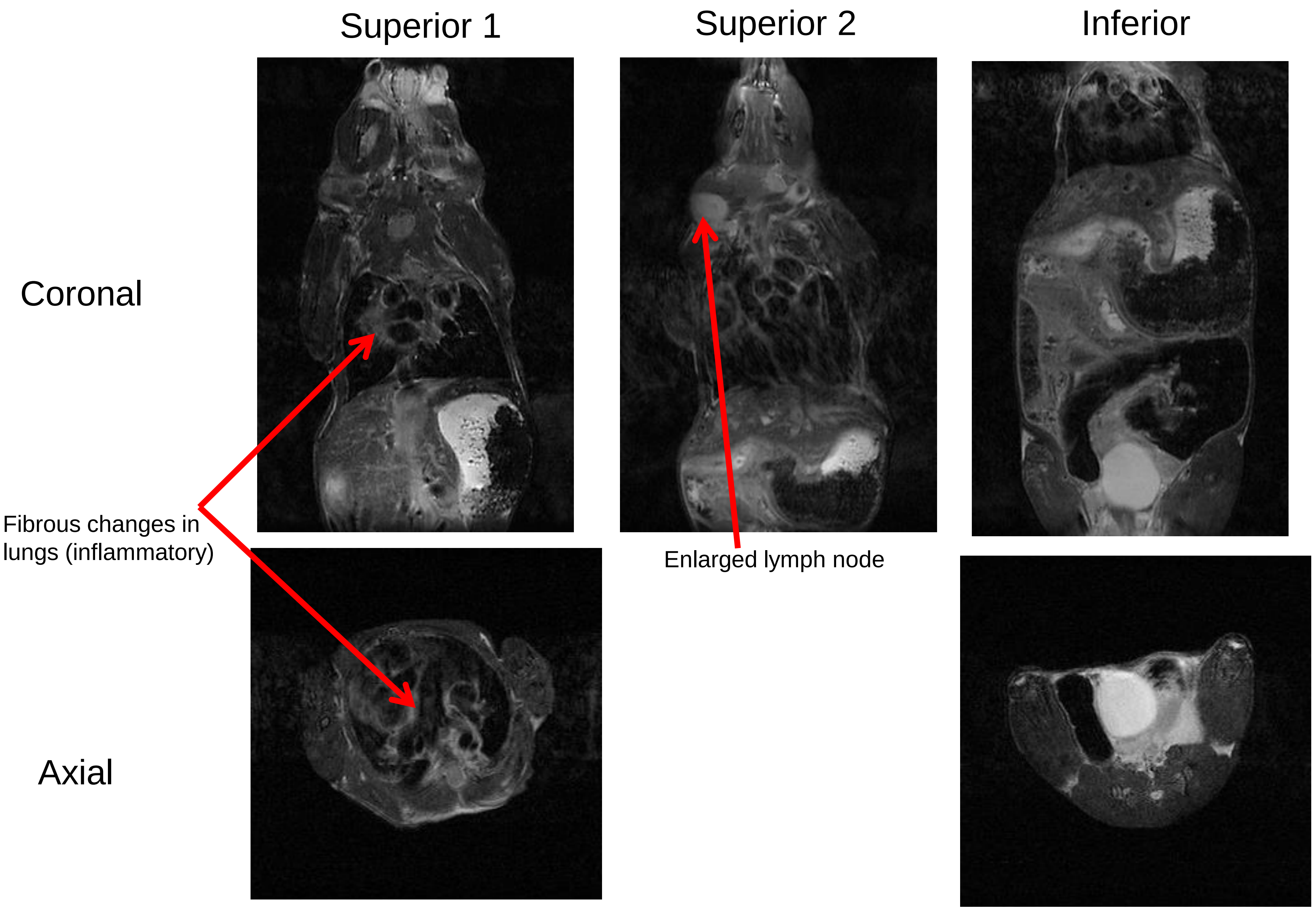

Supp. Fig.2A

LUNG

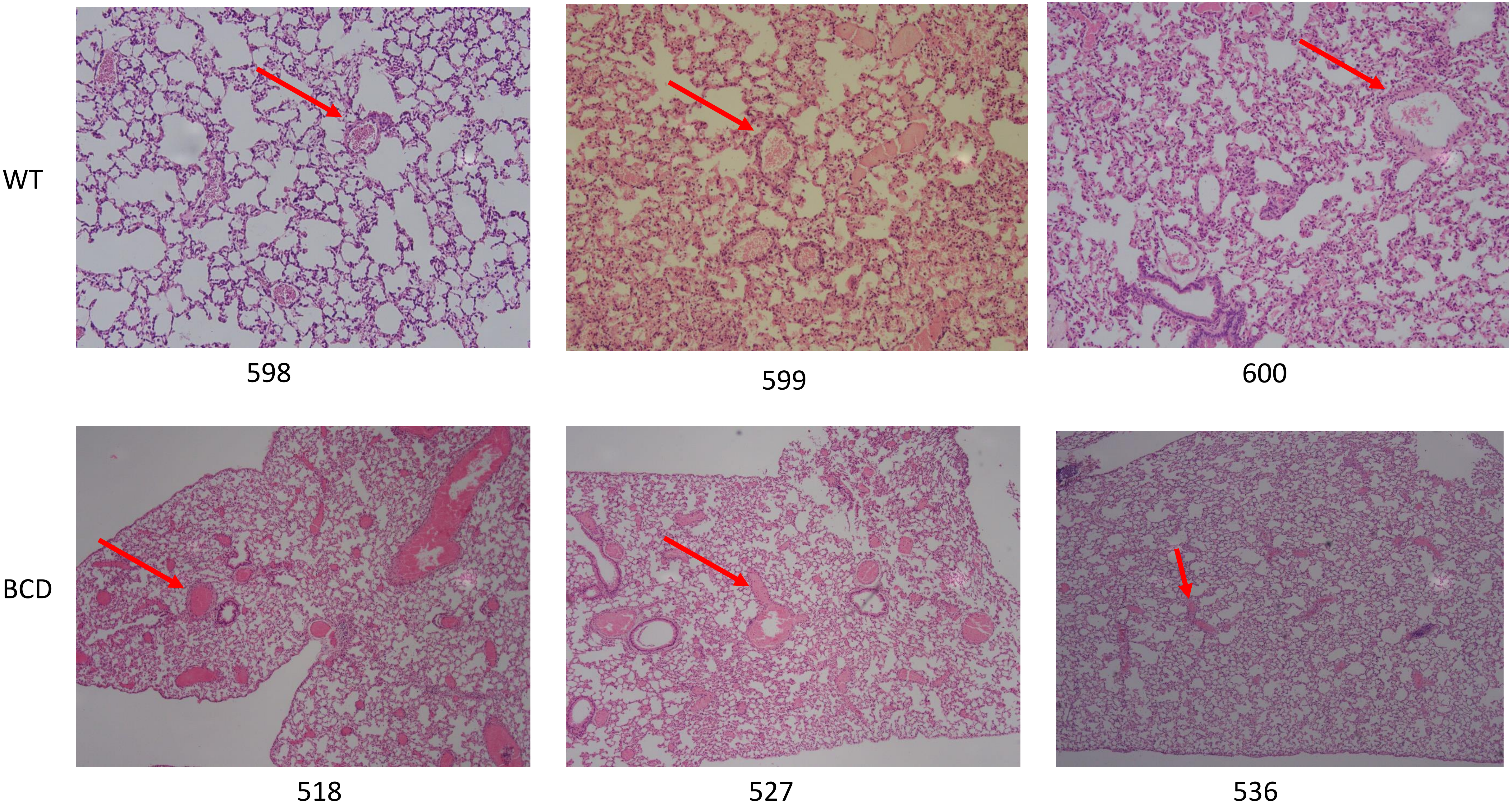

Supp. Fig.2B

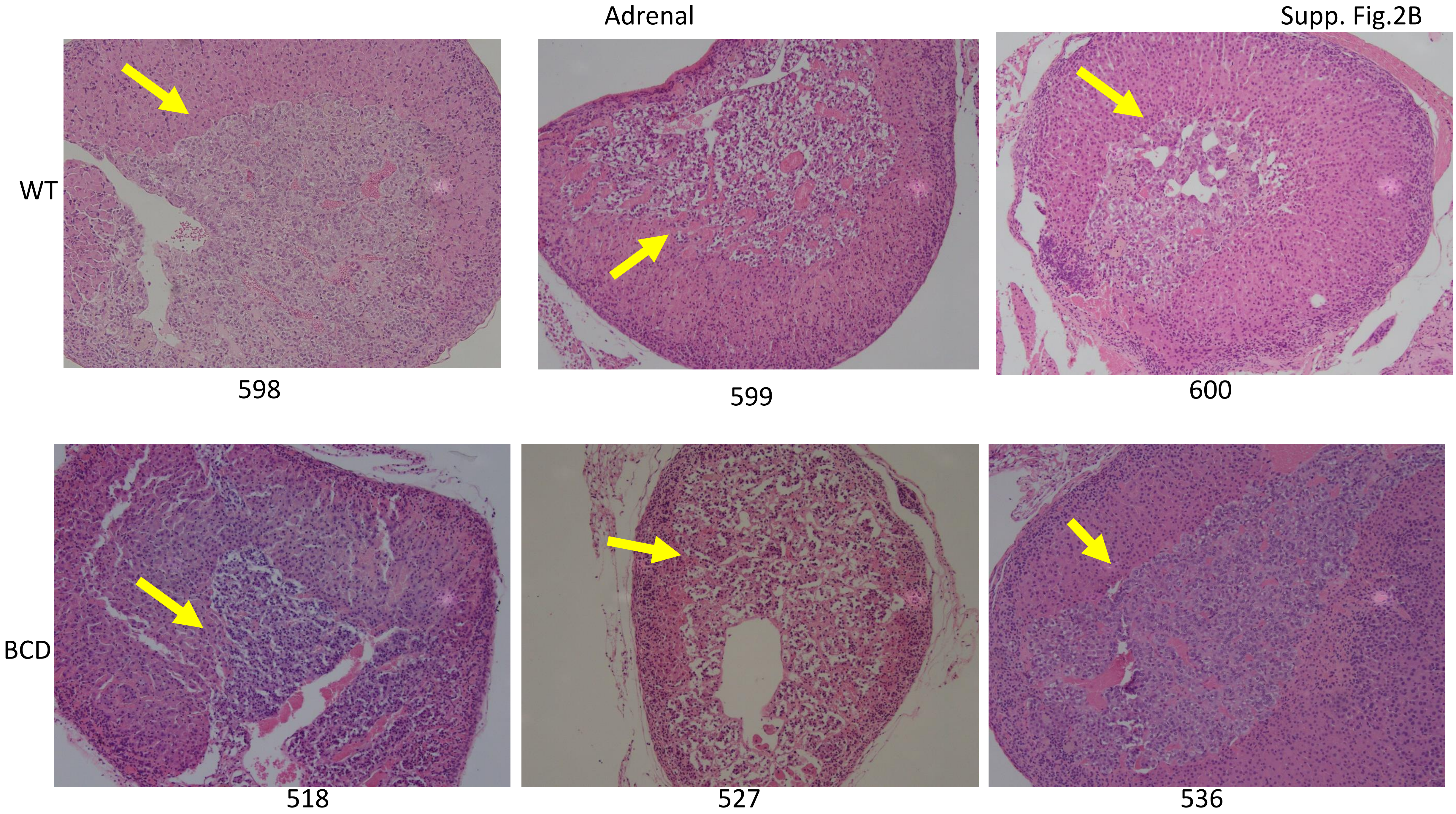

Supp. Fig. 3

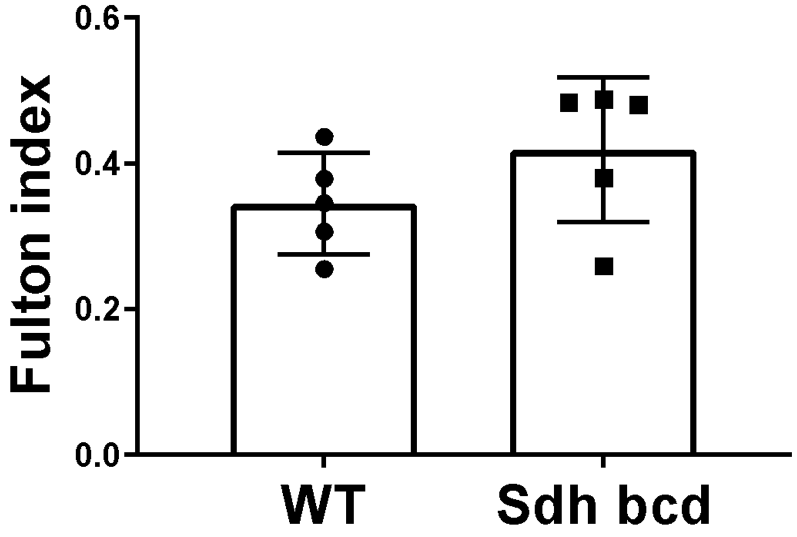
